# Cytosine but not adenine base editor generates mutations in mice

**DOI:** 10.1101/731927

**Authors:** Hye Kyung Lee, Harold E. Smith, Chengyu Liu, Michaela Willi, Lothar Hennighausen

**Affiliations:** Laboratory of Genetics and Physiology, National Institute of Diabetes and Digestive and Kidney Diseases, US National Institutes of Health, Bethesda, Maryland 20892, USA; Genomics Core, National Institute of Diabetes and Digestive and Kidney Diseases, US National Institutes of Health, Bethesda, Maryland 20892, USA; Transgenic Core, National Heart, Lung, and Blood Institute, US National Institutes of Health, Bethesda, Maryland 20892, USA

**Author notes:** These authors contributed equally to this work (Hye Kyung Lee, Michaela Willi, Lothar Hennighausen). Correspondence to: H.K.L, M.W. and L.H.

## Abstract

Deaminase base editing has emerged as a tool to install or correct point mutations in the genomes of living cells in a wide range of organisms and its ultimate success therapeutically depends on its accuracy. Here we have investigated the fidelity of cytosine base editor 4 (BE4) and adenine base editor (ABE) in mouse embryos using unbiased whole genome sequencing of a family-based trio cohort. We demonstrate that BE4-edited mice carry an excess of single-nucleotide variants and deletions compared to ABE-edited mice and controls.

## INTRODUCTION

Deaminase base editing (1,2) directly converts target C•G base pairs to T•A by cytosine base editors (CBE), or target A•T base pairs to G•C by adenine base editors (ABE), without inducing double-stranded DNA breaks (3). Since the majority of known human pathogenic variants are single nucleotide alterations (2,4), base editing has been heralded as a high-fidelity tool to correct single-nucleotide polymorphisms (SNPs) associated with many human disorders. While exceptional precision is paramount in a quest to correct somatic and in particular germline mutations, recent studies have revealed that CBEs can induce bystander mutations, including deletions, in mouse zygotes (5) and plants (6). In contrast, ABE displays a greater fidelity (5,7).

Whole-genome sequencing (WGS) of base-edited rice (8) and mouse embryos (9) revealed that BE3, a commonly used CBE, induces a large number of inadvertent base changes throughout the genome, while ABE displayed high fidelity. In a separate study, WGS of BE3-edited sheep did not reveal an obvious increase of off-target mutations (10). Since BE3 can introduce unwanted indels (1,5) and other undesirable base substitutions in addition to C-to-T conversions (5,7,11), the fourth-generation BE4, containing a second uracil glycosylase inhibitor (UGI) domain and optimized linker architectures, appears to have an increased fidelity *in vitro* (12), in mouse zygotes (5) and in rabbit embryos (13). Off-target effect for BE4 might be expected based on WGS studies that examined off-target mutations introduced by BE3 and ABE (8,9). However, as different sgRNA, editors, and analysis methods might yield different results (8,9), there is a definitive need for investigating *in vivo* genomic effects of BE4 in comparison with ABE. Along those lines, the original study on WGS of CRISPR/Cas9 edited mice suggested the presence of extensive off-target mutations (14), which was, however, likely the result of an imperfect experimental design as pointed out in editorials (15–19). Further WGS investigations by Iyer and colleagues as well as our group (20,21) using trio studies demonstrated that CRISPR/Cas9 does not introduce an excess of off-target mutations. Therefore, it is prudent to examine critical issues of base editing, such as the extent of off-target mutations with a larger number of mice and under several conditions. In this study, we have addressed the question of base editing fidelity and conducted unbiased WGS on a total of 44 BE4- and ABE-edited mice, control mice and their wild-type parents.

## RESULTS & DISCUSSION

To assess on- and off-target fidelities of the advanced BE4 and ABE in mouse embryos, we conducted a family-based trio WGS study (Figure 1A). Fertilized eggs were injected with BE4 or ABE7.10 mRNA together with a sgRNA used by both editors, which permitted a direct comparison. Two-cell stage embryos were implanted into surrogate mothers and 13 ABE-edited and nine CBE-edited founder mice were born, together with 13 non-injected controls (Figure 1B). All founders (Figure 1C and Supplementary Figure 1) were screened by Sanger sequencing to identify mutants and targeted deep-sequencing was performed to determine haplotypes. ABE introduced A-to-G transitions in the target window and, except one, no bystander or proximal off-target mutations were detected in the 33 edited alleles. In contrast, BE4 induced not only the expected C-to-T transitions but also C-to-G and C-to-A conversions in the target site, frequent proximal off-target mutations, and deletions in four of the nine founders (12 out of the 17 edited alleles) (Figure 1C and Supplementary Figure 1).

**Figure 1.**
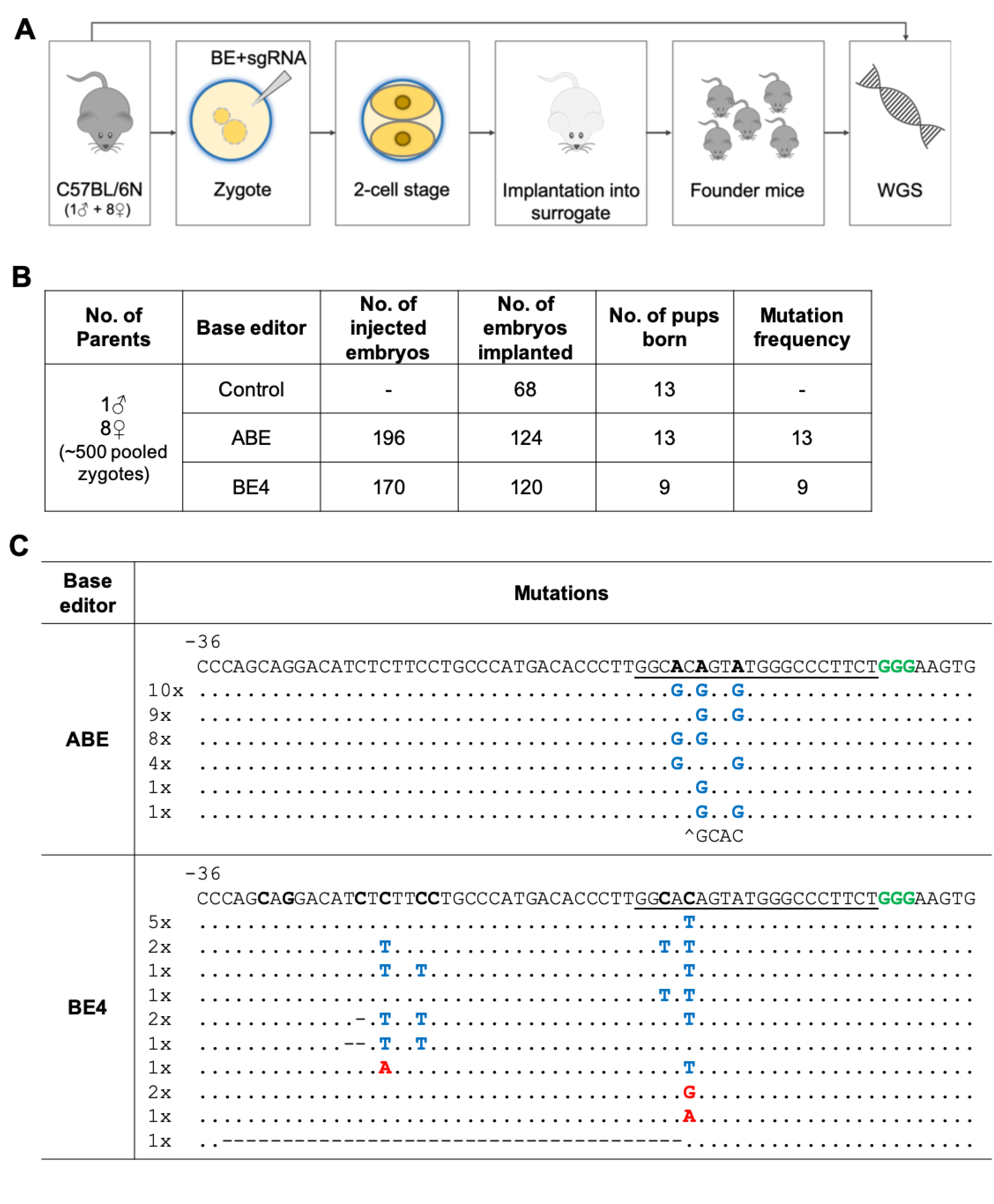
Base-editing by BE4 and ABE in mouse embryos. (**A**) Experimental design of the family-based trio study. (**B**) Summary of data obtained from mouse zygotes co-injected with ABE or BE4 mRNA and a sgRNA. (**C**) Alignment of sequences from founder mice. 33 mutant alleles edited by ABE, and 17 alleles edited by BE4 were analyzed. The sgRNA sequence is underlined and the PAM sites shown in green. The target nucleotides and edited nucleotide are shown in bold black and blue, respectively. Unintended nucleotide substitutions are shown in red. Deletions are marked with a deletion symbol.

Unbiased WGS was performed on the 22 edited mice, 13 controls and nine parents at an average depth of 60x (Figure 1B and Supplementary Table 1). The WGS data were analyzed using GATK with Joint Genotyping and subsequent filtering to identify single nucleotide variants (SNVs) and indels for each individual mouse (Figure 2A). Lumpy with SvTyper was used to identify indels. To explicitly identify *de novo* mutations, the SNVs and indels present in the parents were subtracted from those identified in the progeny (Supplementary Tables 2 and 3). Non-edited control mice had accumulated an average of 132 *de novo* SNVs (Figure 2B and Supplementary Table 2). On average 119 SNVs were detected in ABE-edited mice, comparable to that in controls. In contrast, BE4-edited mice carried on average 221 SNVs, a significant increase (Kruskal-Wallis chi-square: p=0.002), especially C-to-T variants (Figure 2B, Supplementary Figure 2 and Supplementary Table 2).

**Figure 2.**
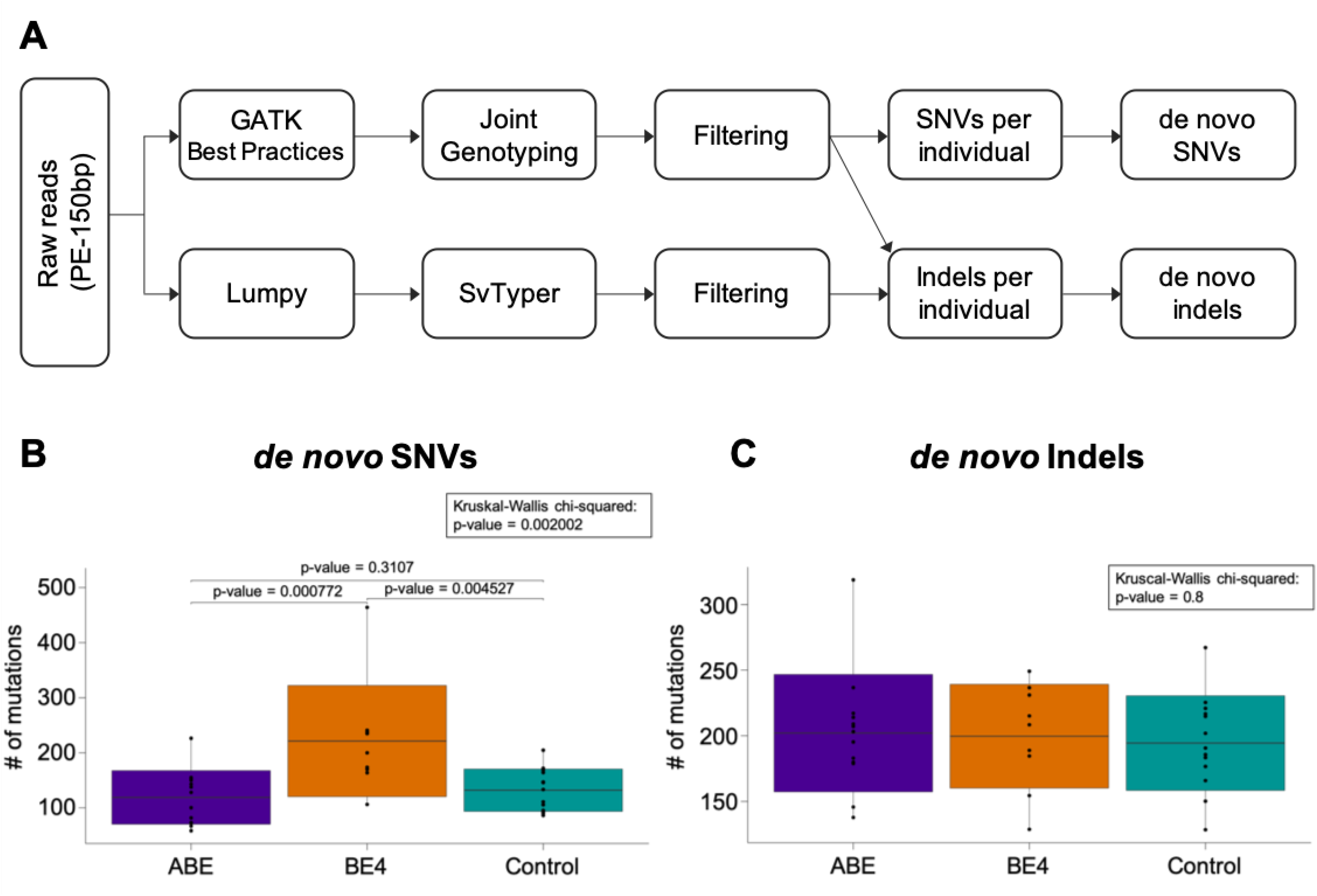
*De novo* mutation frequencies in base-edited and control mice compared to their parents. (**A**) Workflow of whole genome sequencing analysis for SNVs and indels. (**b and c**) Numbers of *de novo* SNVs (**B**) and indels (**C**) identified in mutants and controls. Boxplot: center line, mean; box limits, plus and minus standard deviation; whiskers, minimum and maximum. Wilcoxon rank-sum test and Mann-Whitney-U-Test for the pairwise comparison were used to evaluate the statistical significance of differences between groups.

About 2% of off-target SNVs coincided with the predicted off-target site, CRISPR, suggesting that the majority of mutations were not dependent of the sgRNA and predictable by predicted off-target sites (Supplementary Table 4). The increased off-target editing observed in BE4 but not ABE implies that these mutations were the result of cytosine deaminase AID/APOBEC1 which can induce SNVs in the absence of sgRNAs (1). C-to-T conversions (plus some C-to-A and C-to-G) are overrepresented in *de novo* SNVs observed in BE4-edited mice, consistent with enzymatic activity of BE4. Since four out of the nine BE4-edited mice carried additional deletions proximal to the target region, we analyzed globally their indel frequencies compared with controls (Figure 2C and Supplementary Table 3). The numbers of indels in the BE4- and ABE-edited group showed no differences from the control group (Figure 2C), also not regarding their characteristics (Supplementary Table 5).

Our results confirm and extend previous work that BE3 but not ABE increases off-target SNVs in mouse embryos (9) and rice (8) by analyzing a large data set of nine BE4-edited and 13 ABE-edited mice (a total of 50 mutant alleles). We observed a significant mutation rate with the improved BE4 in mouse embryos. While base-edited mouse embryos (9) acquire off-targets independent on sgRNAs, in base-edited rice off-target mutations can coincide with predicted off target sites (8) suggesting sgRNA dependence. Thus, using the same sgRNA for both BE4 and ABE eliminates latent promiscuous guide targeting as the explanation for off-target mutations in BE4, a unique aspect of our study. We demonstrated an approximately two-fold increase of *de no*vo off-target mutations in BE4-edited embryos, which favorably compares to the more than 20-fold increase observed in BE3-edited mouse embryos (9). However, given the variation between individual mice, our study and that in rice (8) show a statistically significant difference in off-target SNVs. BE4, an advanced version of BE3, contains an additional UGI (uracil DNA glycosylase inhibitor) and a shortened linker, as a means to enhance its specificity (12). Further studies will be needed to validate the higher genome-wide fidelity of BE4. However, BE4 caused inadvertent proximal off-target mutations and deletions in four of the nine founders, which has implications in its use as therapeutic agent. These adverse mutations are likely independent of the sgRNA used as none of the 13 embryos edited with ABE and the same sgRNA displayed any proximal off-target mutations and deletions. It is not clear whether BE3-induced proximal off-target mutations were detected in the two WGS studies (8,9).

In summary, our study emphasizes the high fidelity of ABE as compared to BE4, which also induces increased unwanted base substitutions and deletions in close proximity to the designed target site. Such unwanted mutations are of particular concern when correcting disease-associated SNPs in proteins, as they could adversely alter non-targeted amino acids. Our study also emphasizes the need to monitor off-target mutations in clonal populations, as the analysis of large pools of cells with variable editing, as commonly conducted in *in vitro* experiments (22–24), results in population averaging. Based on our study and previous experiments (5,8,9), ABE appears to be the current choice for base editing because of its fidelity at target sites and throughout the genome. However, caution about the fidelity of deaminase base editors comes from recent studies that demonstrated extensive off-target RNA editing (25,26).

## MATERIALS AND METHODS

### Mice

All animals were housed and handled according to the guidelines of the Animal Care and Use Committee (ACUC) of the NIH (https://oacu.oir.nih.gov) and all animal experiments were approved by the ACUC of National Institute of Diabetes and Digestive and Kidney Diseases (NIDDK, MD) and performed under the NIDDK animal protocol K089-LGP-17. Base edited founder mice were generated using C57BL/6N mice (Charles River Laboratories, MD) by the Transgenic Core of the National Heart, Lung, and Blood Institute (NHLBI, MD).

### CRISPR reagents and microinjection of mouse zygotes

The Wap-E1 sgRNA (GGCACAGTATGGGCCCTTCT) (27), which contains two cytidines and two adenines near the editing window, was designed and synthesized using ThermoFiser’s sgRNA in vitro transcription service. The pCMV-BE4 plasmid (gift from David Liu’s laboratory) and pCMV-ABE7.10 plasmid (Addgene plasmid #102919) were linearized and then their mRNAs were synthesized *in vitro* using the mMESSAGE mMACHINE T7 kit (ThermoFisher Scientific). Mouse zygotes were produced by in vitro fertilization (IVF) using eggs collected from eight superovulated C57BL/6N female mice and sperm collected from one C57BL/6N male (Charles River Laboratories). The ABE and BE4 mRNAs (50ng/ul) were separately microinjected with the sgRNA (20ng/ul) into the cytoplasm of the IVF zygotes. After culturing overnight in M16 medium, those embryos reached 2-cell stage of development were implanted into oviducts of pseudopregnant foster mothers (Swiss Webster, NY). Mice born to the foster mothers were genotyped and subsequently analyzed by whole genome sequencing.

### Genotyping

Genomic DNA was isolated from the tip of tails of 3-week-old mice using Wizard Genomic DNA purification Kit (Promega), amplified by PCR, and followed by Sanger sequencing (Quintarabio, CA). Mutations were identified by PCR amplifying a 599 bp fragment encompassing the target sequence, followed by Sanger sequencing. Library preparation and whole genome sequencing was conducted by the Broad Institute (Cambridge, MA) using Illumina HiSeq X, at a coverage of 60x using 150bp paired-end reads (Supplementary Table 1).

PCR Primers

Wap-S1_F1: GTTGGAACCCATCACAGACAAAGG

Wap-S1_R1: TGTAGAAACAGAGCAGAGAGGTGG

### GATK analysis

WGS (60x) was performed on 44 mice, nine parents (one male and eight females), and their progeny, including 22 founder mice carrying base substitutions at target sites induced by ABE or BE4 using one guide RNA and 13 non-injected control ones. The analysis was performed accordingly to the GATK best practices guidelines (28–30) for germline mutations (version 3.8-0). Quality control and alignment was done by BBmap (31) (version 37.36) and BWA MEM (32) (version 0.7.15), respectively, using the reference genome mm10.

For runtime opitmazation, the aligned BAM files were split up to a chromosome level (for runtime optimization) and reads aligned to different chromosomes were filtered using SAMtools (33) (version 1.5), followed by Picard tools (34) (version 2.9.2) to mark duplicates. The GATK analysis workflow was applied as follows: (i) base recalibration – GATK BaseRecalibrator, AnalyzeCovariates, and PrintReads – using the databases of known polymorphic sites, dbSNP142 and MGPv5 (provided by the high-performance computing team of the NIH (Biowulf)); (ii) variant calling – GATK HaplotypeCaller – with the genotyping mode “discovery”, the “ERC” parameter for creating gvcf and a minimum phred-scaled confidence threshold of 30. The final step included merging the VCF files of each chromosome (GenomeAnalysisTK, GATK).

### GATK SNV analysis

Joint genotyping was applied on on all 44 samples together and hard filters were applied: “QD < 2.0 || FS > 60.0 || MQ < 40.0 || MQRankSum < −12.5 || ReadPosRankSum < −8.0 || SOR > 3 ". The resulting SNVs were additionally filtered by removing those overlapping with repetitive elements (35) (UCSC’s masked repeats plus simple repeats; http://hgdownload.soe.ucsc.edu/goldenPath/mm10/database) and black regions (ENCODE (36); http://mitra.stanford.edu/kundaje/akundaje/release/blacklists/mm10-mouse/). On an individual level, only SNVs with a genotype of 0/1 or 1/1 were kept. Further filtering steps comprised the removal of SNVs with a read depth smaller than 10, an excessive read depth (37) (d+3√d, d = average read depth), an allele frequency less than 10% using a variety of tools (38–40). All SNVs within +/- 5bp of an indel border were removed as likely false-positives.

### Simple GATK indel analysis

Indels identified by GATK where extracted after joint genotyping and subsequently hard filters were applied according to the GATK recommendations: "QD < 2.0 || FS > 200.0 || ReadPosRankSum < −20.0 || SOR > 10.0 ". Indels overlapping with repetitive elements (35) (UCSC’s masked repeats plus simple repeats; http://hgdownload.soe.ucsc.edu/goldenPath/mm10/database) or black regions (ENCODE(36); http://mitra.stanford.edu/kundaje/akundaje/release/blacklists/mm10-mouse/) were removed. The individual samples were filtered keeping only indels with the genotypes of 0/1 and 1/1, removing those with a read depth smaller than 10 as well as sites with an excessive number of reads (37) (d+3√d, d = average read depth). Last, all simple indels that overlap with complex indels identified using LUMPY were excluded. For all those steps a variety of tools (38–40) was applied.

### Complex indel analysis using Lumpy

The analysis of complex indels was done on the same samples using Lumpy (41) according to the guidelines. Mapping was done using BWA MEM (32), with the parameters “--excludeDups --addMateTags --maxSplitCount 2 --minNonOverlap 20” (reference genome mm10), followed by Lumpy (41) using the discordant and split reads as input and genotypes were identified using SVTyper (42). The subsequent filtering steps comprised the selection of indels with a genotype of 0/1 and 1/1 and the removal of indels with a quality smaller than 100 and an excessive read coverage (d+3√d (37), where d is the average read depth) or a SU value (Number of pieces of evidence supporting the variant across all samples) smaller than 5. Indels overlapping with repetitive elements (35) or black regions (36) were excluded.

### Statistical analysis

All statistical analyses were performed with R package 3.3.3 (http://www.R-project.org/). Kruskal-Wallis test was applied using kruskal.test and pairwise comparison was done with a Wilcoxon Rank Sum wilcox.test in R. All values represent means ± S.D.

### Targeted deep-seq

Target sites were amplified from mouse genomic DNA using Phusion polymerase (Thermo Fisher Scientific) and PCR products were prepared as libraries for next-generation sequencing. Pooled PCR amplicons were conducted paired-end sequencing using an Illumina MiSeq (Illumina).

PCR Primers

Wap-S1_F2: AAGACAGGAGGTTTTGAGCAAGGC

Wap-S1_R2: CACCAGTGAAGACAAAGGAGTATGG

### Off-target analysis

Off-target sites were predicted using http://crispor.tefor.net/ (43). The resulting off-target sites were filtered using the same criteria as for SNVs and indels, to consider only those areas of the genome which do not coincide with black regions (36) (ENCODE (36); http://mitra.stanford.edu/kundaje/akundaje/release/blacklists/mm10-mouse) or repetitive elements (35) (UCSC’s masked repeats plus simple repeats; http://hgdownload.soe.ucsc.edu/goldenPath/mm10/database). Mutations, which were present in the population and not only in base-edited mice, but identified at predicted off-target sites, were not considered as a consequence of base editing.

## Data availability

The data are available at SRA under project number PRJNA555149.

## ACKNOWLEDGMENTS

This work utilized the computational resources of the NIH HPC Biowulf cluster (http://hpc.nih.gov). This work was supported by the Intramural Research Programs (IRPs) of National Institute of Diabetes and Digestive and Kidney Diseases (NIDDK) and National Heart, Lung, and Blood Institute (NHLBI).

## FUNDING

This study was supported by the IRPs of NHLBI and NIDDK.

## AUTHOR CONTRIBUTIONS

All authors designed the study. C.L. generated mutant mice. H.K.L. identified and characterized mutant mice. H.K.L. performed experiments and data analysis. M.W. performed computational analysis. H.K.L., M.W. and L.H. supervised the study. H.K.L., M.W. and L.H. wrote the manuscript and all authors approved the final version.

## Competing interests

The authors have not competing interests.

---

**Supplementary Figure 1.**
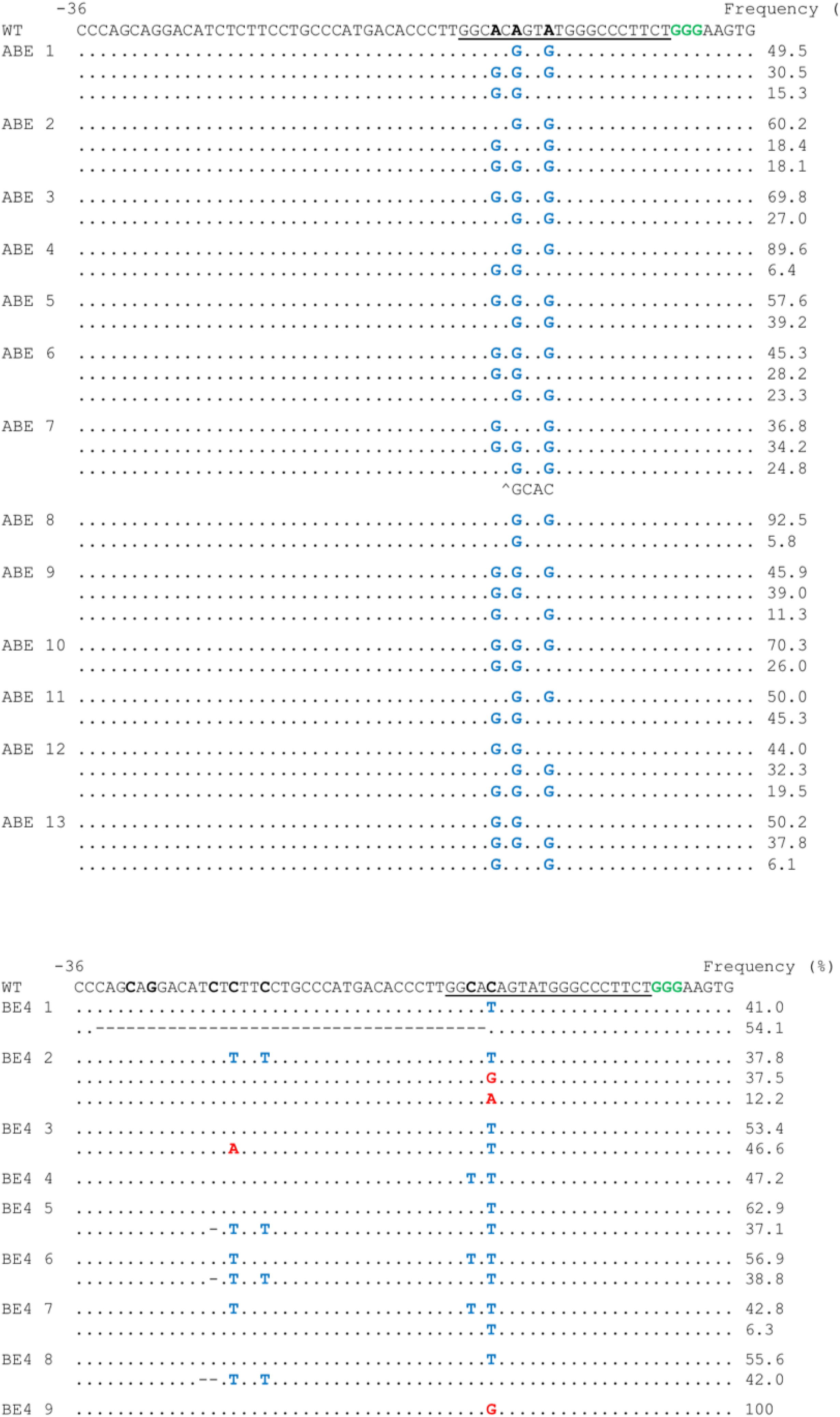
Alignments of mutant sequences from founder mice carrying base-edited mutations. The sgRNA is underlined and the corresponding PAM sequence is shown in green. Within the editing window, the target nucleotides for base editing and nucleotides substituted by base editing are shown in bold black and blue, respectively. Bystanders are highlighted in red and deletions are indicated using a deletion symbol. WT, wild-type.

**Supplementary figure 2.**
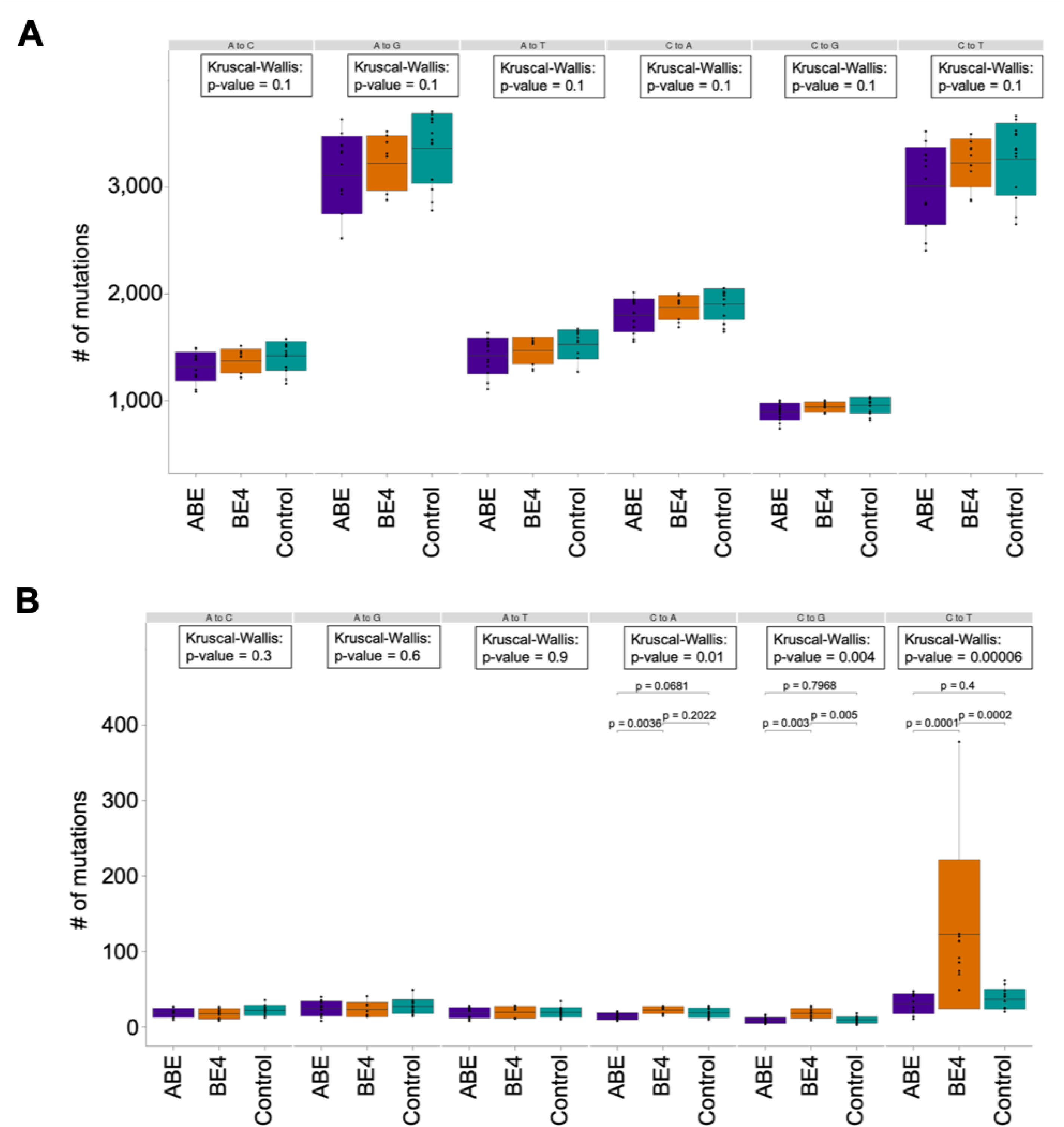
The frequencies of different types of total (A) and *de novo* SNVs (B) in individuals edited by the base editors and in the control group.

**Supplementary Table 1.** Raw read counts of the WGS samples

| Parents |  |  |
| --- | --- | --- |
| Sample | Gender | Raw Read Counts |
| Parent 1 | ♀ | 707,581,699 |
| Parent 2 |  | 868,778,178 |
| Parent 3 |  | 1,072,332,421 |
| Parent 4 |  | 1,162,951,335 |
| Parent 5 |  | 767,582,455 |
| Parent 6 |  | 772,091,887 |
| Parent 7 |  | 823,878,962 |
| Parent 8 |  | 783,498,969 |
| Parent 9 | ♂ | 869,713,827 |

| ABE edited Progeny |  |  |
| --- | --- | --- |
| Sample | Gender | Raw Read Counts |
| ABE 1 | ♀ | 769,235,486 |
| ABE 2 |  | 998,329,339 |
| ABE 3 |  | 1,033,798,370 |
| ABE 4 |  | 808,263,614 |
| ABE 5 |  | 651,033,776 |
| ABE 6 | ♂ | 839,652,961 |
| ABE 7 |  | 1,137,405,701 |
| ABE 8 |  | 735,637,641 |
| ABE 9 |  | 919,901,911 |
| ABE 10 |  | 881,365,098 |
| ABE 11 |  | 686,200,552 |
| ABE 12 | N/A | 1,141,706,029 |
| ABE 13 |  | 874,997,141 |

| Control Progeny |  |  |
| --- | --- | --- |
| Sample | Gender | Raw Read Counts |
| Control 1 | ♀ | 945,102,788 |
| Control 2 |  | 945,102,788 |
| Control 3 |  | 945,102,788 |
| Control 4 |  | 914,556,327 |
| Control 5 | ♂ | 767,715,791 |
| Control 6 |  | 935,639,522 |
| Control 7 |  | 651,111,242 |
| Control 8 |  | 669,733,921 |
| Control 9 |  | 671,835,774 |
| Control 10 |  | 768,856,803 |
| Control 11 |  | 875,378,967 |
| Control 12 |  | 667,712,201 |
| Control 13 |  | 842,759,887 |

| BE4 edited Progeny |  |  |
| --- | --- | --- |
| Sample | Gender | Raw Read Counts |
| BE4 1 | ♀ | 721,100,551 |
| BE4 2 |  | 792,525,208 |
| BE4 3 |  | 764,834,081 |
| BE4 4 | ♂ | 952,740,731 |
| BE4 5 |  | 743,607,260 |
| BE4 6 |  | 905,084,101 |
| BE4 7 |  | 798,497,190 |
| BE4 8 |  | 686,497,626 |
| BE4 9 |  | 1,013,557,705 |

**Supplementary Table 2.**
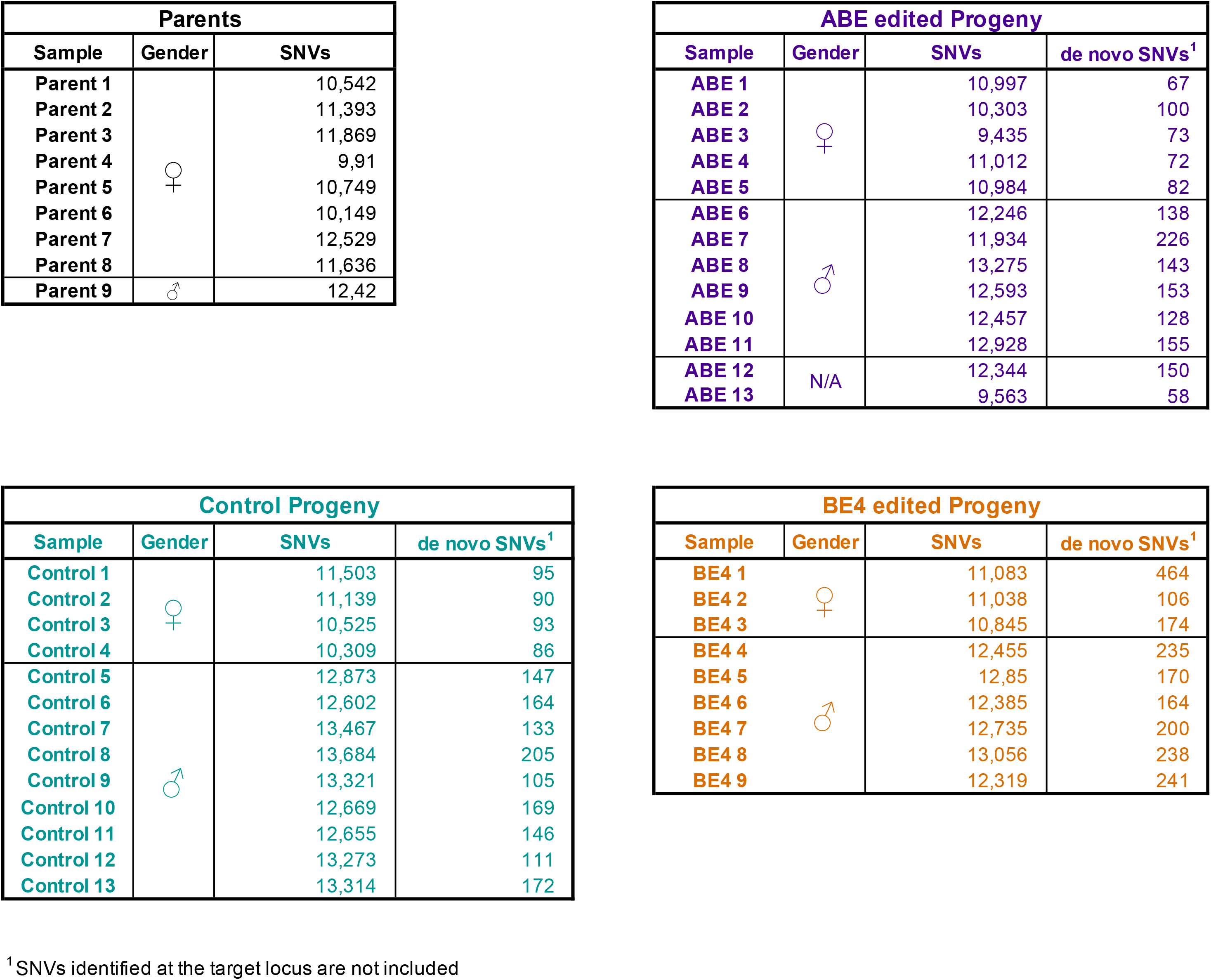
SNVs identified in each sample and *de novo* SNVs of the progeny

**Supplementary Table 3.**
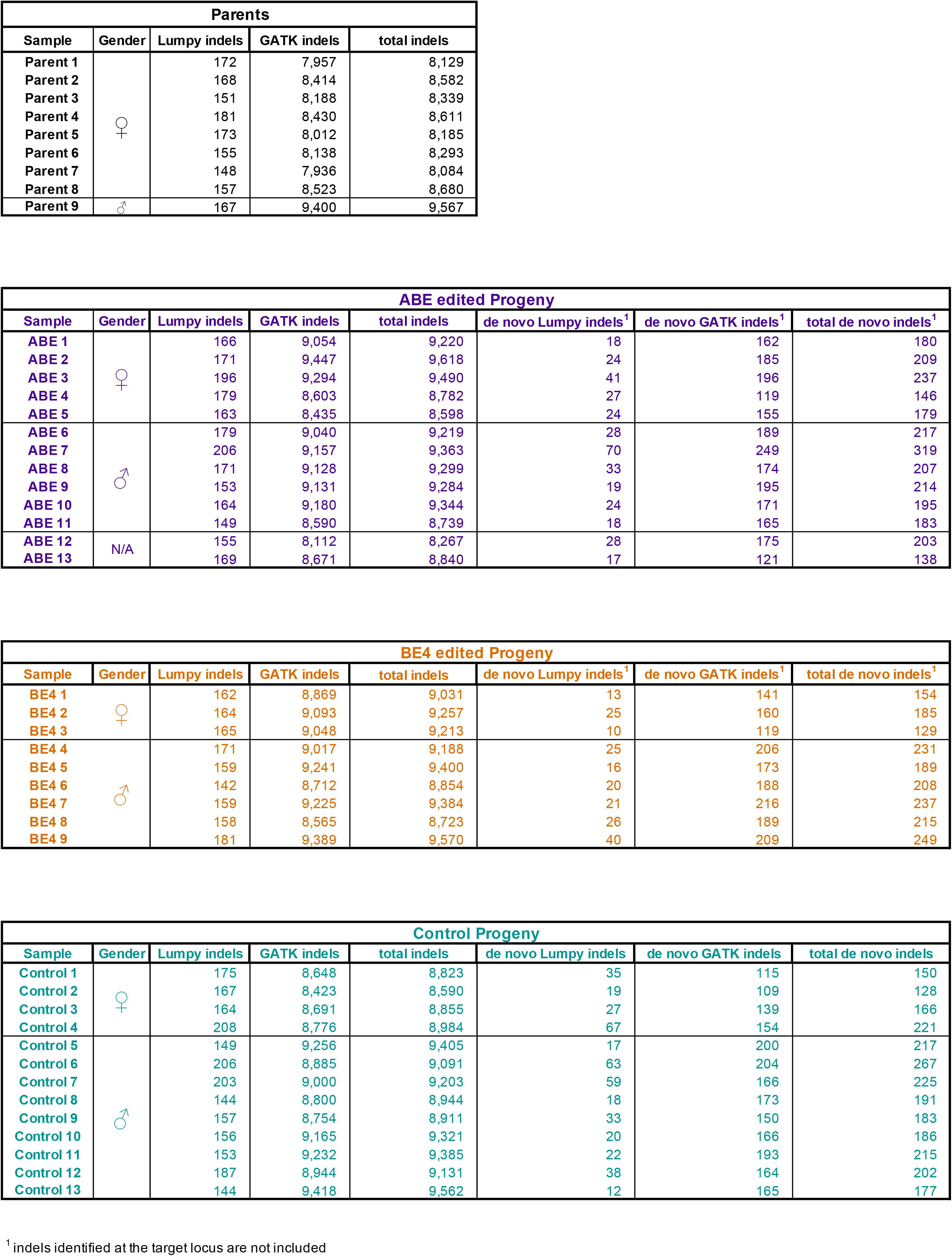
Indels identified in each sample and *de novo* indels of the progeny

**Supplementary Table 4.** Location of the four predicted off-target sites overlapping with *de novo* SNVs. The samples harboring the same SNV are grouped into overlapping groups.

| off-target coordinate |  |  | SNVs groups overlapping with the off-target site |  |  |  |
| --- | --- | --- | --- | --- | --- | --- |
| chr | start | stop | Overlap 1 | Overlap 2 | Overlap 3 | Overlap 4 |
| chr11 | 60377453 | 60377475 | ABE 5 | BE4 7<br>BE4 8 | BE4 1<br>BE4 3<br>BE4 7 | BE4 8 |
| chr13 | 114169949 | 114169971 | ABE 1 |  |  |  |
| chr9 | 40294521 | 40294543 | ABE 2<br>ABE 4 | ABE 3<br>ABE 4<br>ABE 6<br>ABE 7<br>ABE 11 |  |  |
| chrX | 17247050 | 17247072 | ABE 1 | ABE 9 | BE4 4 |  |

**Supplementary Table 5:** Characterization of *de novo* indels.

|  | GATK indels |  |  |
| --- | --- | --- | --- |
|  | ABE edited<br>mean (+/- s.d.) | BE4 edited<br>mean (+/- s.d.) | Control<br>mean (+/- s.d.) |
| deletions | 24% (+/- 6.1) | 27% (+/- 6.5) | 29% (+/- 4.1) |
| insertions | 76% (+/- 6.1) | 73% (+/- 6.5) | 71% (+/- 4.1) |
| avg. deletion size (bp) | 2.7 (+/- 0.8) | 2.9 (+/- 0.7) | 2.9 (+/- 0.5) |
| avg. insertion size (bp) | 15.2 (+/- 3.4) | 13.3 (+/- 2.5) | 12 (+/- 3.1) |

|  | LUMPY indels |  |  |
| --- | --- | --- | --- |
|  | ABE edited<br>mean (+/- s.d.) | BE4 edited<br>mean (+/- s.d.) | Control<br>mean (+/- s.d.) |
| deletions | 64.1% (+/- 13.2) | 57.6% (+/- 10.1) | 64.4% (+/- 18.6) |
| duplications | 0.7% (+/- 1.7) | 0% (+/- 0) | 0.6% (+/- 1.5) |
| inversion | 0.6% (+/- 2.3) | 0% (+/- 0) | 0.8% (+/- 1.7) |
| complex | 34.6% (+/- 11.5) | 42.4% (+/- 10.1) | 34.1% (+/- 18.4) |
| avg. deletion size (bp) | 672 (+/- 332) | 1,001 (+/- 843) | 2,218 (+/- 4,343) |
| avg. duplication size (bp) | 123,900 (+/- 27,788) | N/A | 52,376 (+/- 73,358) |
| avg. inversion size (bp) | 115 (+/- 0) | N/A | 12,516 (+/- 17,452) |

